# Widespread 3-Base Periodicity in Complex DNA Mixtures and an Alignment-Free Algorithm to Uncover its Source

**DOI:** 10.64898/2026.09.26.754620

**Authors:** Noam Hecht, Nicolas Duek, Yuval Dotan, Saharon Rosset, Viviane Slon, Muli Safra

## Abstract

**Background:** Protein-coding DNA exhibits characteristic three-base periodicity. In unaligned DNA libraries, however, this signal should cancel if fragment boundaries are uniformly distributed across codon phases. Yet, we find that in multiple metagenomic libraries, base frequencies depended systematically on distance from fragment boundaries, producing a period-3 pattern. We investigated the origin of this unexpected signal.

**Results:** The signal was observed in modern metagenomic datasets and widely across ancient-DNA datasets. We developed a new alignment-free algorithm to infer latent triplet-phase structure from nucleotide composition. The inferred shifts were strongly nonuniform at fragment boundaries, explaining the read-coordinate periodicity. We initially suspected a protocol artifact. To test an alternative, we derived a probabilistic framework linking nucleotide-dependent boundary selection to codon-phase frequencies. The framework showed that, because nucleotide composition differs among codon positions, preferential breakage at particular nucleotides biases the phases represented at fragment boundaries. Simulations confirmed this mechanism: purine-associated fragmentation produced strong period-3 coherence in a coding-dense bacterial genome, whereas no comparable genome-wide effect appeared in the human reference genome.

**Conclusions:** The observed read-coordinate three-base periodicity does not require an intrinsic phase bias in the source DNA. Instead, sequence-dependent fragment formation or recovery can couple boundaries to codon phase. Purine-associated post-mortem fragmentation plausibly explains the widespread phenomenon in ancient-DNA libraries, although PCR- or library-specific mechanisms may contribute to it in modern datasets. Our alignment-free algorithm provides a practical method for estimating and normalizing phase shifts before searching for positional nucleotide patterns, preventing boundary-associated bias from being mistaken for an intrinsic biological pattern.

## 1 Background

Sequencing libraries consist of DNA fragments whose ends define a coordinate system that is generally unrelated to the biological coordinate system of the source genome. DNA molecules may already be fragmented in the original material or may be fragmented during extraction and library preparation; both biological degradation and laboratory fragmentation can produce non-random, sequence-dependent fragment boundaries [1–3]. The resulting molecules are then processed, potentially amplified by PCR, and sequenced. A priori, there is no reason for the nucleotide composition at an internal position of a read to depend systematically on its distance from a fragment boundary. Indeed, if fragment formation, library recovery, and amplification do not preferentially select particular sequence phases, then fragments beginning at many different genomic positions should be represented in all possible phases. Any compositional pattern tied to genomic coordinates should consequently lose its fixed relationship to the read boundaries after the fragments are pooled.

Thus, even before considering periodicity, a reproducible dependence of nucleotide frequencies on distance from a read boundary is unexpected. A dependence specifically on this distance modulo three is more surprising still: it suggests that the fragment-defined coordinate system has somehow remained coupled to the codon coordinate system of protein-coding DNA. While examining complex DNA libraries, we observed precisely such a phenomenon. For many exact read lengths *ℓ*, nucleotide frequencies varied systematically with distance from the fragment boundary, separating into three recurring profiles according to position modulo three. Closely related profiles reappeared for read lengths separated by multiples of three. Importantly, this structure was visible directly in the sequenced reads, before alignment, assembly, taxonomic assignment, or coding-sequence annotation. At first sight, this observation raises a potentially serious methodological concern. Library construction and PCR amplification are complex, sequence-dependent processes, and the frequencies of reads in a sequenced library need not faithfully reproduce the relative abundance of molecules in the original DNA mixture [4–6]. A strong read-coordinate period-3 pattern could therefore indicate that amplification, recovery, or another stage of the experimental protocol has preferentially retained particular classes of fragments. If its source were unknown, such a distortion would be difficult to distinguish from a genuine biological signal and could complicate downstream comparisons of composition or abundance.

We first encountered the phenomenon prominently in ancient environmental DNA datasets. Ancient DNA provides an informative setting in which to study fragment-boundary effects because its molecules are typically short and have undergone extensive post-mortem fragmentation and chemical alteration [3, 7, 8]. Sedimentary ancient DNA (sedaDNA) is additionally taxonomically complex and can contain mixtures of microbial, plant, animal, and fungal DNA. Such material can recover information about past ecosystems and hominin presence even in the absence of diagnostic macrofossils or well-preserved skeletal remains [9–11]. Its analysis commonly relies on short-read mapping or *k*-mer matching against reference databases [12–14]. Classifier performance can depend strongly on read properties and database composition, and reference-derived sequence similarity can produce false taxonomic assignments in ancient environmental metagenomes [15, 16]. As shown below, however, the read-coordinate phenomenon is neither confined to a single ancient DNA study nor exclusive to ancient DNA. The analyses developed below trace this pattern to unequal representation of the three inferred codon-phase shifts as a function of distance from the fragment boundary. This mechanism helps resolve the broader methodological concern: the observed periodicity need not reflect an arbitrary discrepancy between the sequenced library and the original DNA mixture. Rather, it can arise from a specific coupling between sequence-dependent boundary formation or recovery and the three-base composition of coding regions. Because this coupling can be detected statistically and its phase imbalance can be normalized, it also provides a practical way to prevent the resulting read-coordinate pattern from confounding downstream analyses.

An introductory example is provided by A2424, a library generated from ancient cave sediment [9]. Among unique sequences of length 59 bp, nucleotide frequencies show a strong and regular dependence on position modulo three (Figure 1).

**Figure 1:**
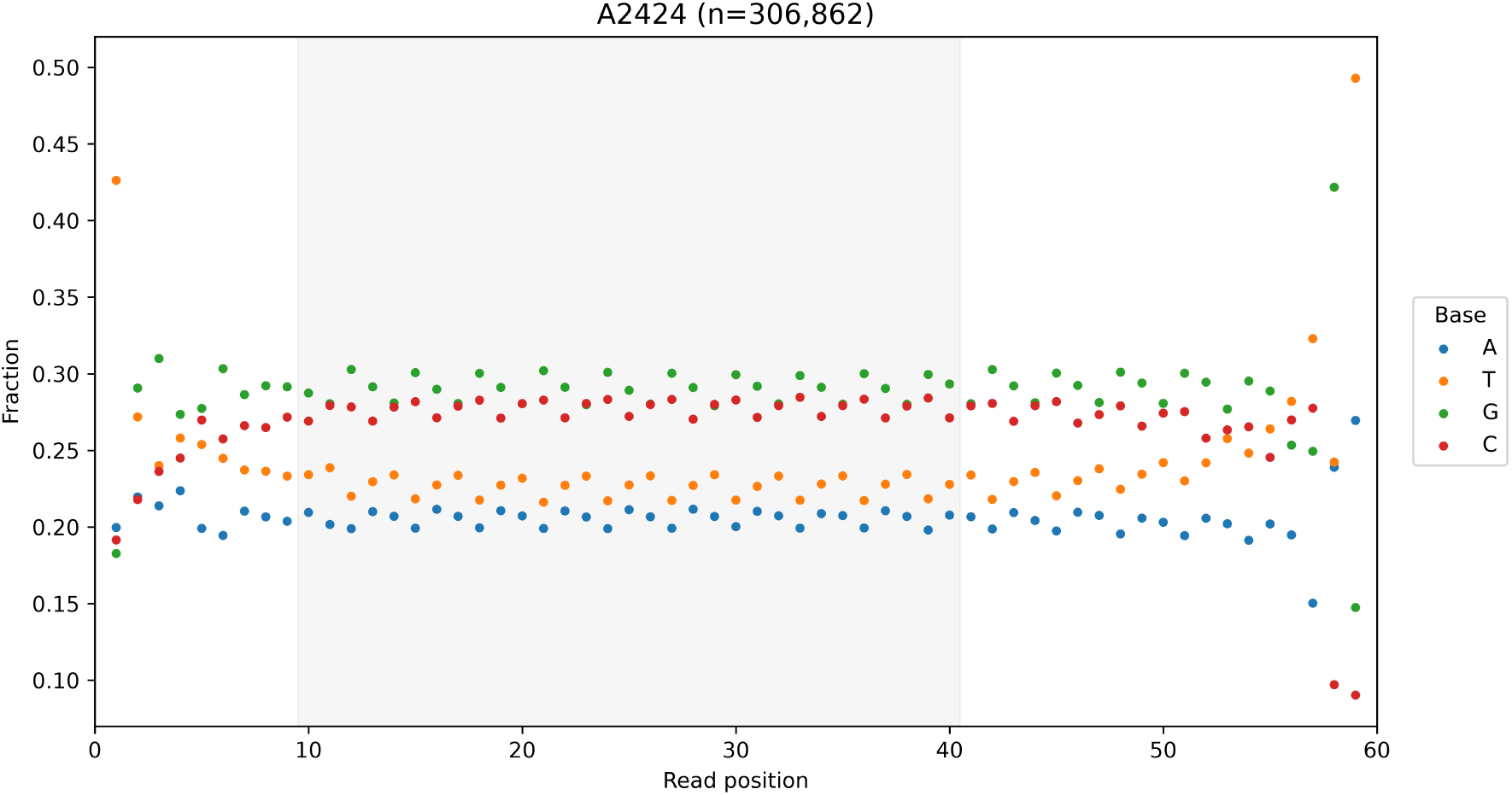
Introductory example of the mod-3 phenomenon. A2424 is a DNA library generated from ancient cave sediment. Shown here are the unique sequences of length 59 base pairs (bp) (*n* = 306,862), which provide an example of the read-coordinate pattern. Nucleotide frequencies depend markedly on read position *i* mod 3, producing three recurring position classes across read coordinates. The gray shaded region marks the regression analysis window, read positions 10–40 inclusive.

Three-base periodicity is a classical property of protein-coding DNA because nucleotide composition and selective constraint differ among the three codon positions [17, 18]. Bacterial genomes are often coding-dense, whereas only a small fraction of the human nuclear genome is protein-coding [19–22]. The presence of coding sequence alone, however, cannot explain a period-3 pattern that remains locked to the ends of sequenced fragments. If fragments begin without preference among the three coding phases, then aligning those fragments by their sequenced ends mixes the three phases and strongly attenuates the aggregate period-3 signal. The observation of a coherent period-3 pattern in read coordinates therefore implies not only an underlying triplet-specific nucleotide composition, but also a mechanism that represents the three possible phases unequally relative to fragment boundaries.

### 1.1 Conceptual framework

Let *p*_*r*_(*b*) denote the probability of nucleotide *b* at coding position class *r ∈ {*0, 1, 2}, and let *s* denote the coding phase at which a fragment begins. At zero-based read coordinate *i*,

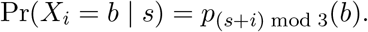

If fragment-start phases occur with probabilities *π*_0_, *π*_1_, *π*_2_, then

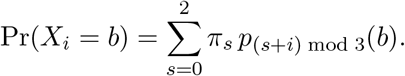

When *π*_0_ = *π*_1_ = *π*_2_ = 1*/*3, the aggregate probability is independent of *i* mod 3. A read-coordinate period-3 signal therefore requires both an underlying triplet-specific nucleotide composition and nonuniform representation of fragment phases.

Sequence-dependent fragment formation or recovery provides a plausible route to phase imbalance. Ancient DNA fragmentation and multiple library-preparation steps can be sequence-dependent [1, 3, 4, 7, 23, 24]. If boundary-associated sequence contexts differ among codon positions, such biases can preferentially represent particular phases at read starts or ends. The coding phase at the terminal base is (*s* + *l −* 1) mod 3; consequently, stable biases acting at both boundaries predict related phase mixtures for read lengths sharing the same value of *ℓ* mod 3.

We investigated this framework first through exact-length composition profiles, period-3 regression, and phase redistribution. We also developed frequency-domain, latent-state, coding-frame-normalization, and fragmentation-simulation workflows.

## 2 Results

### 2.1 Read lengths with the same remainder modulo three show similar nucleotide-frequency profiles

Read-position single-nucleotide composition profiles showed a regular three-position pattern in the A2424 ancient cave-sediment library. Within the internal read positions, the overall nucleotide composition was comparatively stable, whereas the relative fraction of each nucleotide oscillated with period three (Figure 2a). Averaging read positions by their residue class modulo three compressed the trajectories into three-position modulo-3 profiles (Figure 2b).

**Figure 2:**
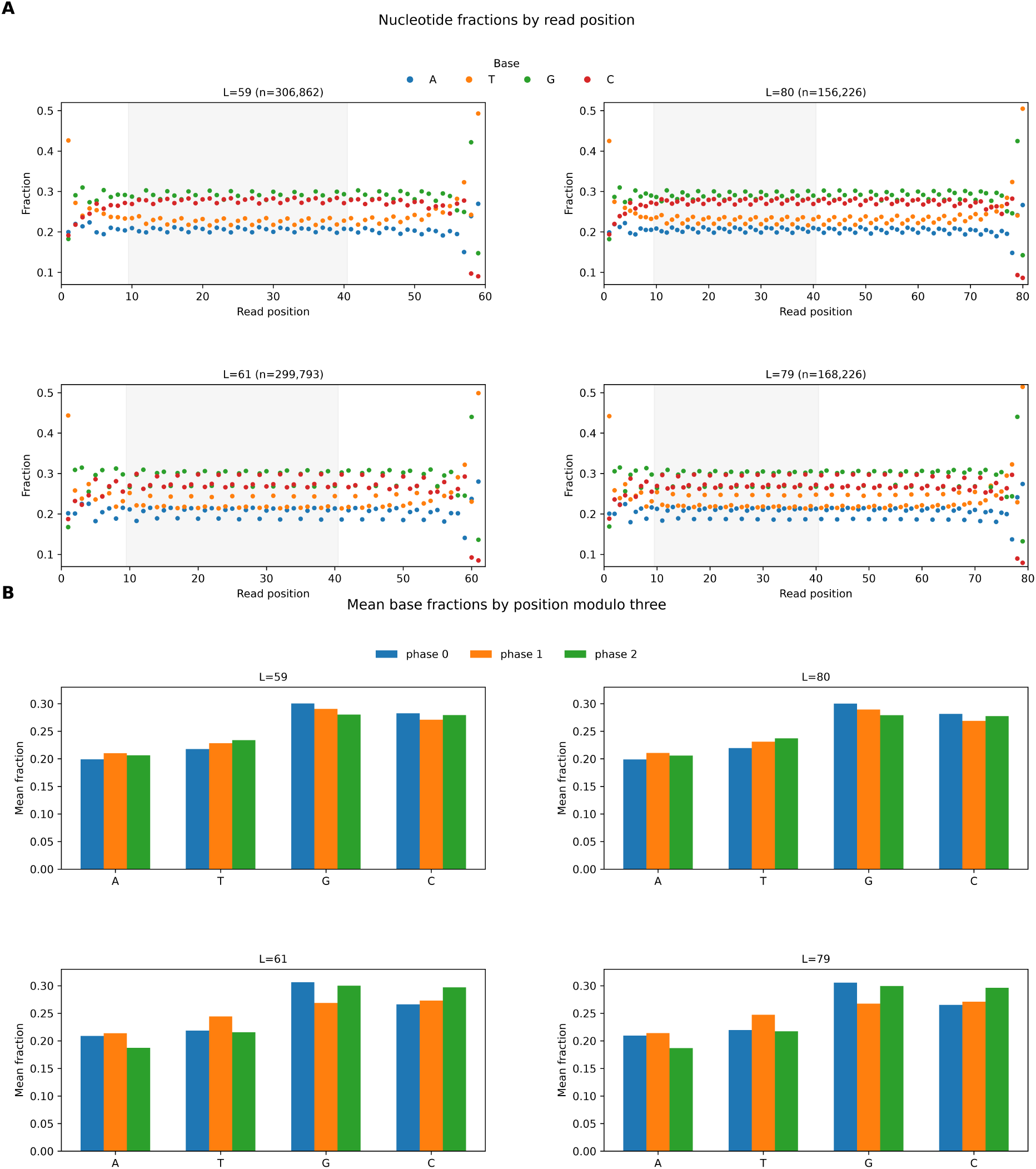
Read lengths in the same modulo-3 class have similar nucleotide-frequency profiles. **A**, Nucleotide-frequency profiles by read position. Lengths 59 and 80 belong to the same modulo-3 class (59 ≡80 (mod 3)), as do lengths 61 and 79 (61 ≡79 (mod 3)). The gray shaded regions mark the regression analysis window, read positions 10–40 inclusive. **B**, Mean nucleotide fractions within read positions 10–40 after grouping positions by their value modulo three. Read lengths within the same modulo-3 class show closely similar modulo-3 profiles, whereas the two modulo-3 classes show different profiles.

The profiles for 59-bp and 80-bp reads were closely matched. Because

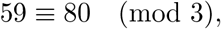

these lengths belong to the same modulo-3 equivalence class. A similar correspondence was observed for 61-bp and 79-bp reads, which likewise satisfy

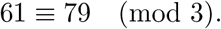

These two pairs illustrate the broader modulo-3 relationship: read lengths belonging to the same modulo-3 class show similar modulo-3 profiles, whereas lengths from different modulo-3 classes show different profiles (Figure 2a,b). This behavior is consistent with a stable relationship between read boundaries and an underlying three-position composition pattern. Strong period-3 fits were observed across the analyzed read lengths; detailed regression statistics are given below for the 59-bp population.

For the 59-bp read population (*n* = 306,862 unique sequences), period-3 regression over read positions 10–40 explained 93.8%, 94.3%, 98.2%, and 94.8% of the positional variance in A, T, G, and C frequencies, respectively. The corresponding fitted period-3 amplitudes were 0.00650, 0.00950, 0.01171, and 0.00702. All four nucleotide-specific period-3 components were significant after Holm correction (*P*_adj_ *<* 0.001). Thus, the high *R*^2^ values reflected a highly regular but modest-amplitude oscillation in nucleotide composition, with the largest periodic component observed for G.

### 2.2 Phase redistribution disrupts read-coordinate period-3 coherence

To test whether the observed oscillation depends on coherent alignment of the three-base pattern across reads, we redistributed A2424 length-59 sequences among the three possible retained phase offsets. Each sequence was shortened by exactly two nucleotides to a final length of 57 bp. The two removed nucleotides were distributed between the sequenced 5^*′*^/3^*′*^ ends using one of three trimming configurations: 0/2, 1/1, or 2/0. These configurations therefore shift the retained 5^*′*^ boundary by zero, one, or two nucleotides, respectively, and represent the three possible read-coordinate phase offsets.

For the phase-redistribution control, each unique original sequence was assigned to one of these three trimming configurations. Exact duplicate sequences received the same assignment within a randomization replicate. Four independent randomization replicates were generated using different seeds; the seed determines the assignment of unique sequences among the three trimming configurations but does not otherwise alter the sequences or the downstream analysis.

As noted above, in the original length-59 population (*n* = 306,862 unique sequences; Figure 3A), the period-3 amplitudes were 0.00650, 0.00950, 0.01171, and 0.00702 for A, T, G, and C, respectively, with corresponding *R*^2^ values of 0.938, 0.943, 0.982, and 0.948. Random phase redistribution strongly attenuated this structure across all four independent replicates. In seed 20, which retained the largest residual period-3 component among the four randomization replicates (Figure 3B), the corresponding amplitudes were only 0.000532, 0.000406, 0.000596, and 0.000398. Even in this conservative replicate, the amplitudes were therefore reduced by approximately 92–95% relative to the original population. The corresponding *R*^2^ values were 0.166, 0.054, 0.158, and 0.062, and none of the four nucleotide-specific period-3 components was significant after Holm correction.

**Figure 3:**
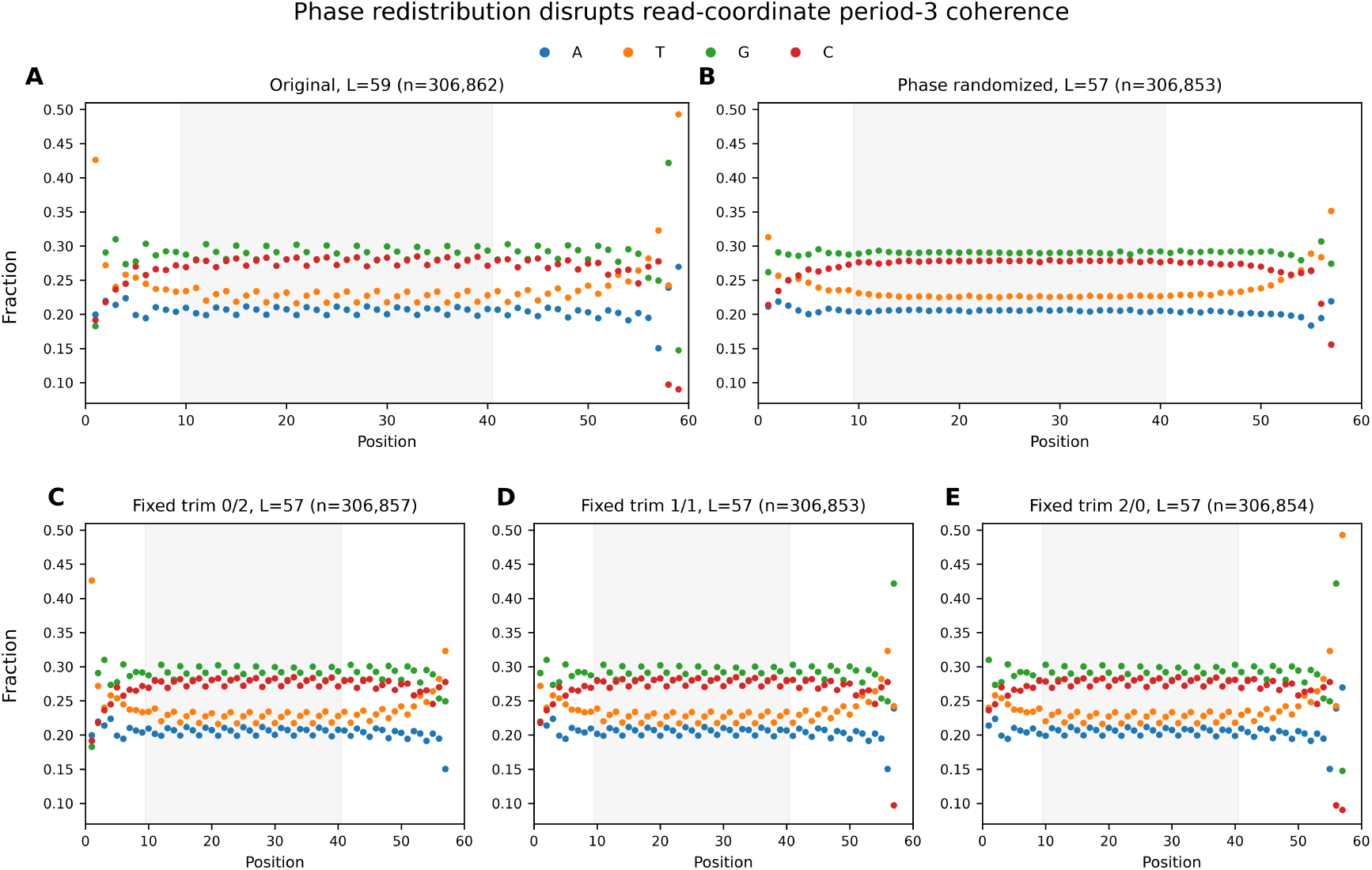
Period-3 structure depends on phase coherence across reads. Nucleotide-frequency profiles by read position for A2424. The gray shaded region marks the regression analysis window, read positions 10–40 inclusive. Small differences in unique-sequence counts among the trimmed datasets result from exact-sequence collapsing after trimming. **A**, Original 59-bp sequences. **B**, Phase-randomized 57-bp sequences (seed 20), generated using the three possible trimming configurations (0/2, 1/1, and 2/0). **C–E**, Fixed-trim controls using 0/2, 1/1, and 2/0 trimming, respectively. Random phase redistribution strongly attenuated the period-3 structure, whereas fixed trimming preserved it.

To verify that this attenuation resulted from mixing phase offsets rather than from shortening the reads, we also generated three phase-preserving controls in which the same trimming configuration was applied to every sequence (Figure 3C–E). All three fixed-trim controls therefore had the same final length of 57 bp as the randomized sequences. Fixed trimming preserved the oscillation amplitude and high period-3 *R*^2^, while successive one-nucleotide displacements of the retained 5^*′*^ boundary shifted the fitted phase by approximately 120^*°*^. Thus, shortening the reads by two nucleotides was not sufficient to abolish the signal; attenuation occurred when sequences with different retained phase offsets were mixed within the same population.

### 2.3 The oscillatory signal is spectrally localized at frequency 1/3

We next represented the nucleotide-frequency profiles in the frequency domain using a discrete Fourier transform (DFT). We analyzed positions 10–39, giving a 30-position window in which period three corresponds exactly to Fourier bin *q* = 10, or frequency *f* = 1*/*3.

The original A2424 length-59 reads showed a pronounced spectral peak at *f* = 1*/*3 for all four nucleotides (Figure 4A). The normalized DFT magnitudes at this frequency were 0.00331, 0.00475, 0.00585, and 0.00352 for A, T, G, and C, respectively. The corresponding spectral variance fractions were approximately 0.948, 0.943, 0.985, and 0.945.

**Figure 4:**
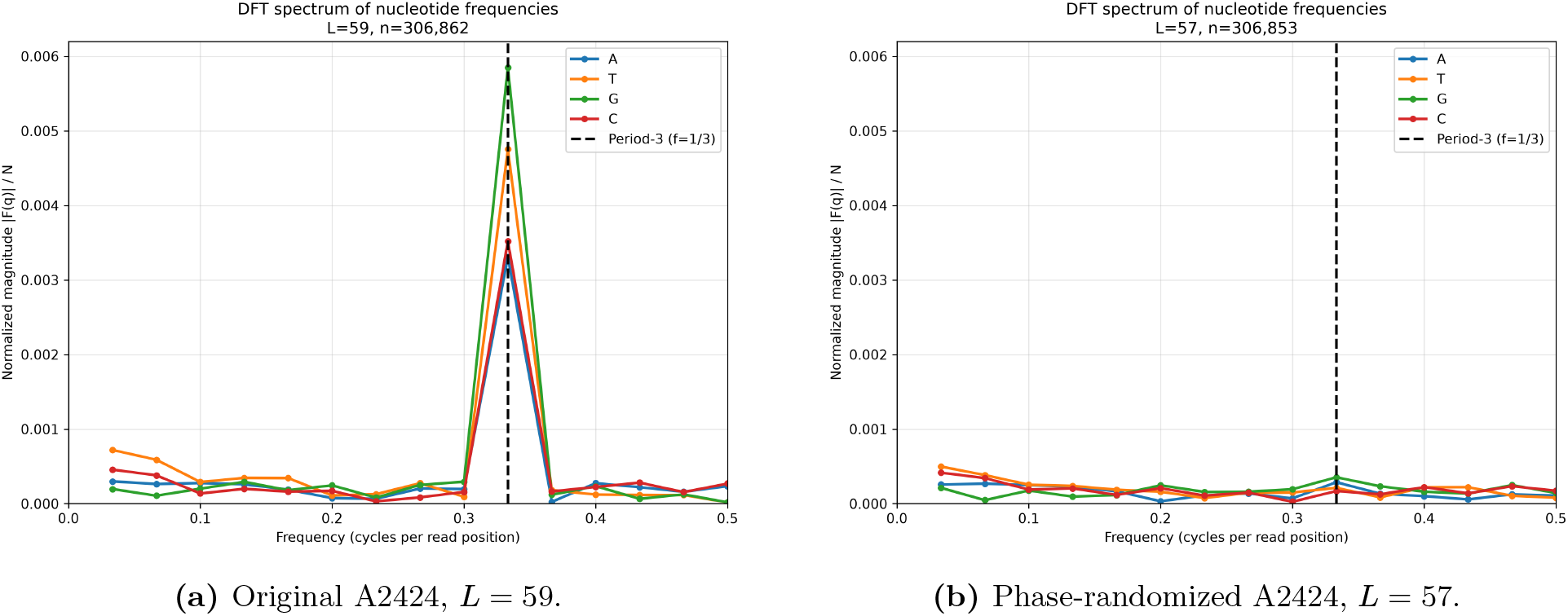
Frequency-domain representation of period-3 structure before and after phase redistribution. **A**, Original A2424 (*L* = 59). **B**, Phase-randomized A2424 (*L* = 57; seed 20), showing the replicate with the largest residual *f* = 1*/*3 component among the four randomization replicates. DFT spectra were calculated over read positions 10–39, placing the period-3 frequency exactly at *f* = 1*/*3 (dashed line).

Phase redistribution markedly attenuated the *f* = 1*/*3 component. Even in seed 20, which had the largest residual period-3 component among the four randomization replicates examined (Figure 4B), the normalized G magnitude decreased from 0.00585 to 0.000354, a 93.9% reduction. The other randomized replicates showed still smaller residual components.

The DFT and sine–cosine regression describe the same harmonic projection and are therefore not independent statistical tests. Rather, the DFT demonstrates that the observed oscillatory structure is specifically localized at the expected period-3 frequency.

### 2.4 Period-3 structure occurs across independent ancient DNA studies

To determine whether read-coordinate period-3 structure was specific to A2424, we extended the analysis to environmental ancient-DNA libraries from AncientMetagenomeDir [25]. For each library, exact read-length classes containing at least 40,000 observed reads were analyzed separately. At each eligible length, the four nucleotide-specific *R*^2^ values were averaged, and the read length with the largest mean *R*^2^ was selected to provide one descriptive period-3 score per library.

Period-3 structure was observed across libraries from numerous independent projects (Figure 5). Several projects contained libraries with selected mean *R*^2^ values approaching one, whereas other projects showed weaker or more heterogeneous structure. Substantial variation was also observed among libraries within individual projects. Thus, strong read-coordinate period-3 structure was not restricted to A2424 or to a single ancient-DNA study, although its magnitude varied considerably across libraries and projects.

**Figure 5:**
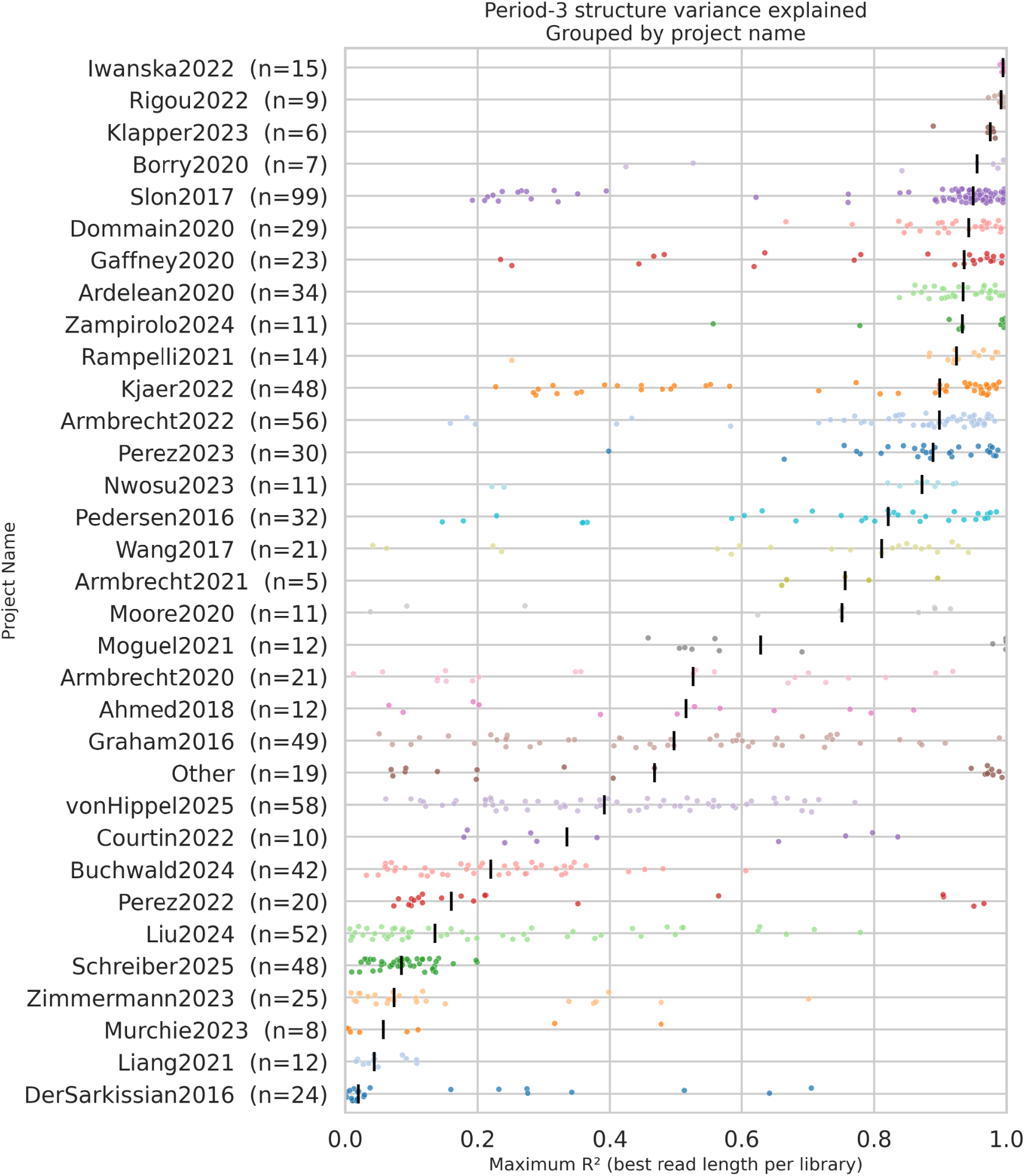
Period-3 structure across independent ancient-DNA projects. Each point represents one library and shows the largest mean period-3 *R*^2^ across its eligible exact read-length classes, where the mean was calculated across A, T, G, and C. Libraries are grouped by project, and black markers indicate project medians. The selected values are used as descriptive library-level summaries rather than as multiplicity-adjusted significance statistics.

### 2.5 Modern metagenomes also show strong period-3 structure

To determine whether read-coordinate period-3 structure was restricted to ancient-DNA libraries, we examined three modern metagenomic datasets representing distinct environments: agricultural soil [26], a Tara Oceans marine metagenome [27], and a human-associated metagenome [28]. Because these longer-read datasets showed stronger terminal composition effects, period-3 regression was evaluated over a common internal window of positions 30–60.

All three datasets showed strong period-3 fits within this internal window (Table 1), with nucleotide-specific *R*^2^ values ranging from 0.935 to 0.995. These examples demonstrate that strong read-coordinate period-3 structure is not restricted to ancient-DNA libraries and can also occur in modern metagenomic sequencing data. Because these datasets were selected examples rather than a systematic survey of modern metagenomes, they do not provide an estimate of the prevalence of the phenomenon in modern sequencing datasets.

**Table 1:** Period-3 structure in selected modern metagenomic datasets. Three datasets representing agricultural soil, marine, and human-associated metagenomes were examined as examples of period-3 structure in modern sequencing libraries.

| Accession | Environment | Length | Window | Reads | $R_A^2$ | $R_T^2$ | $R_G^2$ | $R_C^2$ |
| --- | --- | --- | --- | --- | --- | --- | --- | --- |
| SRR25158531 | Agricultural soil | 150 | 30–60 | 70,893,337 | 0.992 | 0.988 | 0.992 | 0.995 |
| ERR599146 | Marine | 100 | 30–60 | 5,903,542 | 0.942 | 0.987 | 0.993 | 0.974 |
| DRR014179 | Human-associated | 110 | 30–60 | 123,207 | 0.944 | 0.943 | 0.951 | 0.935 |

### 2.6 Coding-frame normalization strengthens triplet-specific composition

To examine the relationship between the read-coordinate signal and coding sequence structure, we identified A2424 reads that mapped unambiguously to annotated coding sequences in a concatenated reference comprising the Unified Human Gastrointestinal Genome (UHGG) catalog v2.0 and the human GRCh38 reference genome. We retained reads mapping in the sense orientation and entirely within a single coding sequence, and normalized them to the annotated coding frame using the GFF phase field. From the same accepted source reads, we constructed a matched control retaining coding orientation but without phase-dependent frame normalization.

For the 11,094 matched reads yielding normalized length 75, the unnormalized control already showed substantial period-3 structure. The fitted amplitudes for A, T, G, and C were 0.0108, 0.0366,0.0512, and 0.0169, respectively. Coding-frame normalization of the identical source reads increased these amplitudes to 0.0146, 0.0935, 0.1233, and 0.0297, corresponding to 1.34-, 2.56-, 2.41-, and 1.75-fold increases.

Thus, coding-derived reads contain substantial three-base compositional structure even before coding-frame normalization, while normalization to the annotated coding phase markedly increases its amplitude. This result supports a connection between the observed periodicity and codon-position-dependent sequence composition, but does not establish that coding reads account for all of the whole-library signal.

### 2.7 Purine-associated fragmentation generates period-3 coherence in a genome-dependent manner

We tested whether nucleotide-dependent fragmentation can generate read-coordinate period-3 structure by simulating fixed-length fragments from the *Pseudomonas aeruginosa* and human reference genomes. Candidate fragments were sampled globally across each reference and assigned either molecular orientation with equal probability. Fragment acceptance depended on purine identity at the read-oriented boundaries. Seven bias conditions were tested, from unbiased random fragmentation to complete purine-associated selection at both boundaries (1.00/1.00), with ten independent simulations of 100,000 reads per condition.

In *P. aeruginosa*, increasing 5^*′*^ purine-associated fragmentation produced a pronounced increase in period-3 structure (Figure 6A). Mean *R*^2^ across A, T, G, and C increased from approximately 0.13 under random fragmentation to 0.82 at 0.70/0.50 and 0.93 at 1.00/0.50. Imposing complete purine selection at both boundaries produced a slightly lower mean *R*^2^ of approximately 0.87, indicating that the effects of the two boundary constraints are not simply additive.

**Figure 6:**
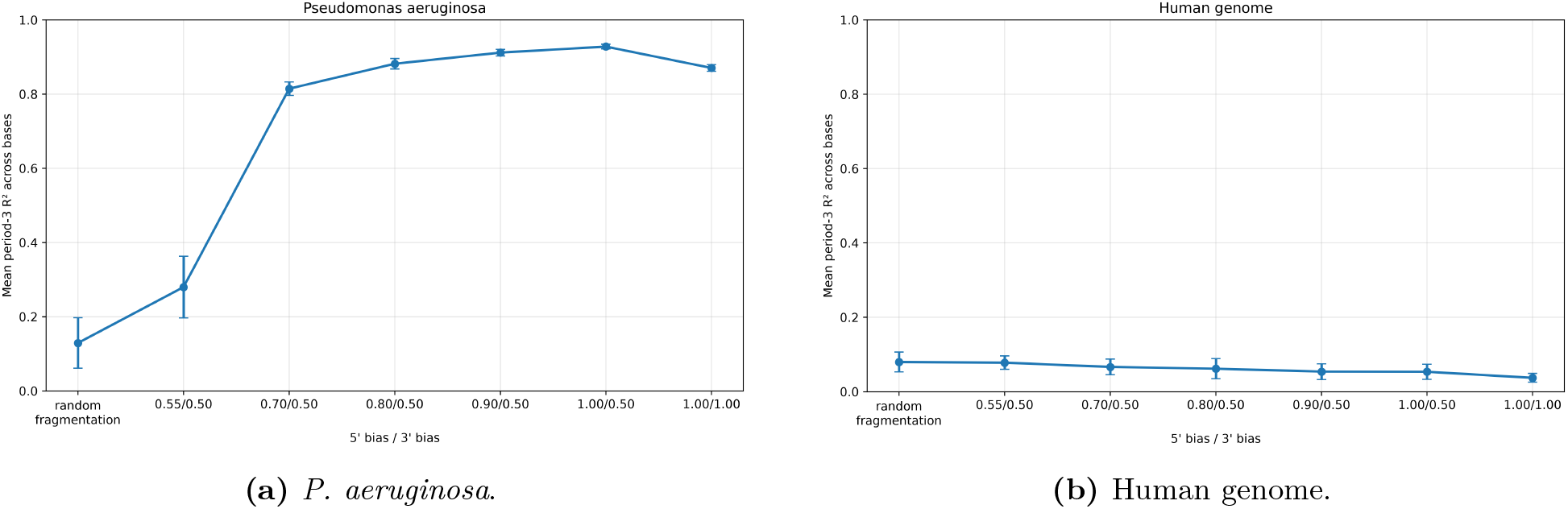
Effect of purine-associated fragmentation on period-3 structure. Each point is the mean period-3 *R*^2^ across A, T, G, and C, summarized across ten independent simulations; error bars show 95% *t* confidence intervals across replicate-level means. The random-fragmentation control was generated without nucleotide-dependent boundary selection, such that fragment boundaries reflected the underlying reference sequence. The remaining conditions imposed the indicated 5^*′*^/3^*′*^ purine-associated fragmentation biases.

The human genome showed no comparable emergence of coherent period-3 structure: mean *R*^2^ remained low across the tested conditions (Figure 6B). Thus, purine-associated fragmentation was sufficient to generate strong period-3 coherence in the bacterial reference but not genome-wide in the human reference. These simulations indicate that fragmentation bias alone is not sufficient in arbitrary sequence and instead interacts with the underlying organization of the source genome.

### 2.8 A six-state latent model recovers triplet-specific sequence composition

We used a six-state hard expectation–maximization (EM) model to characterize triplet-specific nucleotide composition while allowing each sequence to adopt one of three cyclic offsets in either orientation. Across A2424 read lengths 59–61, the inferred 3 *×* 4 nucleotide-probability matrix was highly stable (Figure 7A).

**Figure 7:**
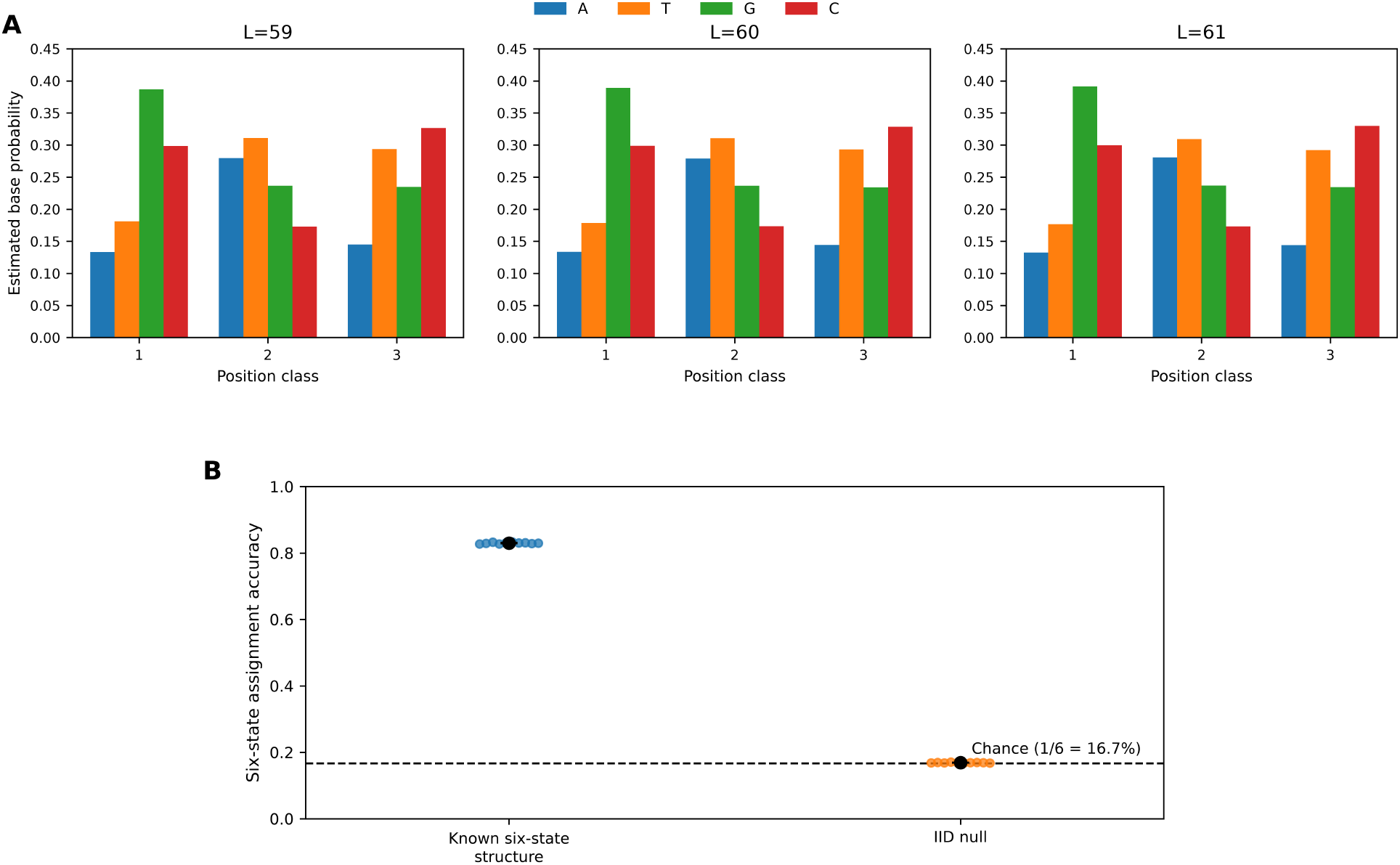
Six-state latent modeling recovers reproducible triplet-specific structure. **A**, Estimated nucleotide probabilities across the three latent position classes for A2424 reads of lengths 59–61 bp, showing stable triplet-specific composition across neighboring read lengths. **B**, Six-state assignment accuracy for ten synthetic datasets with known six-state structure and ten composition-matched IID-null datasets. Points represent independent datasets; black markers indicate the mean, and error bars show 95% t-based confidence intervals. The dashed line indicates chance-level accuracy (1*/*6 = 16.7%).

Random redistribution of read phase strongly attenuated the aggregate read-coordinate signal but did not eliminate the triplet-specific matrix recovered by the latent-state model. This distinction is expected because the six-state model explicitly allows the phase of each sequence to vary: phase redistribution removes coherence among reads while preserving within-sequence triplet composition that can be recovered by latent phase reassignment.

We validated model recovery using ten independently generated synthetic datasets of 60,000 reads from a known triplet-specific nucleotide-probability matrix with balanced latent phase/orientation states. The recovered matrices had a mean root-mean-square error of 0.0036 relative to the generating parameters, and latent six-state assignments were recovered with 83.0% mean accuracy after accounting for model label symmetries. In composition-matched independent and identically distributed (IID) null datasets, nucleotides were sampled independently from the same overall base-frequency distribution at every sequence position, thereby removing position-class dependence. Assignment accuracy was 16.9%, close to the six-state chance expectation of 16.7% (Figure 7B).

All 200 positive-control restarts converged to assignment-stable solutions, compared with 197 of 200 IID-null restarts. Multiple initializations remained necessary because both empirical and synthetic analyses contained inferior local optima; the highest-scoring converged solution among 20 independent restarts was therefore retained for each analysis.

## 3 Discussion

The central observation of this study is that nucleotide composition in unaligned metagenomic reads can display a meaningful bias as a function of the distance from the read boundary, producing a recurrent period-3 pattern. This differs from the classical observation of three-base periodicity in protein-coding DNA: codon-position-dependent composition should not remain visible when fragment boundaries are uniformly distributed across codon phases. Within the framework considered here, the signal arises when underlying triplet-specific composition is coupled to unequal representation of phases relative to read boundaries.

Several analyses support this interpretation. Read lengths in the same modulo-3 class exhibited similar phase bias. Random redistribution of retained read offsets strongly attenuated the aggregate periodicity, whereas fixed-trim controls preserved it. These controls show that the observed oscillation depends on read-coordinate coherence; they do not, by themselves, identify its biological or technical source. The stronger triplet-specific composition obtained after normalizing a mapped subset to annotated coding frames independently supports a contribution from coding sequence. Because that subset is reference-selected, however, it does not establish the fraction of the whole-library signal attributable to coding-derived reads. Fourier analysis provides another representation of the same period-3 component rather than an independent statistical test.

An initially concerning possibility was that the boundary-associated signal reflected an unrecognized distortion introduced during library preparation or amplification. Sequence-dependent fragmentation provides a mechanistically specific alternative that does not require such a failure. The relevant probabilistic distinction is between the phase distribution at candidate genomic positions and the phase distribution conditional on a fragment being formed and recovered. Because nucleotide composition differs among codon positions, a preference for particular boundary nucleotides can make the latter distribution nonuniform. Conditioning on fragment formation or recovery can therefore reweight the represented phases according to their sequence-dependent recovery probabilities. Thus, a biochemical preference that does not explicitly recognize codon position can nevertheless produce a phase-dependent sampling pattern. Note, however, that this explanation does not establish that the recovered library is an unbiased representation of the original DNA mixture, nor does it exclude additional amplification or library-preparation effects.

The simulations demonstrate this mechanism’s sufficiency in a controlled setting. Purine-associated boundary selection generated pronounced period-3 coherence in the *Pseudomonas aeruginosa* reference, whereas no comparable genome-wide response was observed in the sparsely coding human reference. Indeed, the effect of boundary selection depends on the sequence to which it is applied. It is consistent with a contribution from coding-rich sequence, but does not isolate coding density from other differences between the two genomes. Moreover, the simulations select fixed-length candidate fragments using specified boundary preferences; they do not reproduce the full sequence of degradation, extraction, end processing, amplification, and sequencing. They therefore establish a plausible mechanism rather than the quantitative cause of the empirical signal.

Purine-associated fragmentation is a possible contributor in ancient DNA, given the documented association between purines and ancient-DNA breakpoints [7, 8, 23]. Nevertheless, occurrence in an ancient environmental library does not by itself establish that the signal-bearing molecules are ancient or that their boundaries arose through post-mortem degradation. The observation of this phenomenon in modern DNA shows that ancient-DNA damage is not the only cause of the signal. Several processes may produce similar read-coordinate patterns, and the differences among libraries could reflect source composition, fragmentation, recovery, or subsequent processing. Distinguishing these possibilities will require comparisons of mapped breakpoint contexts and annotated coding phases, together with protocol-matched controls. Interpretation must also distinguish observed read ends from molecular fragment ends: an unmerged read can end before the underlying fragment does, and trimming can alter its apparent boundaries.

The alignment-free six-state model can be used for estimating latent triplet-specific nucleotide composition while allowing individual sequences to differ in phase and orientation. This analysis provides an important distinction between the observed read-coordinate pattern and the underlying triplet-specific composition. Read lengths belonging to different modulo-3 classes can exhibit different nucleotide-frequency profiles in read coordinates, yet the inferred 3 *×* 4 nucleotide-probability matrix was highly stable across neighboring read lengths spanning the three modulo-3 classes. This is consistent with the length-dependent profiles representing different boundary-relative phases of a shared underlying triplet-specific nucleotide composition, rather than fundamentally different triplet compositions at different read lengths. The result does not imply that all sequences share the same taxonomic origin, function, or biological history; rather, it indicates that a common triplet-specific nucleotide composition can be recovered despite differences in how that structure is phased relative to read boundaries.

This interpretation also explains why random phase redistribution strongly attenuates the aggregate read-coordinate oscillation while leaving triplet-specific composition recoverable by the latent-state model. Phase redistribution disrupts coherence among reads, whereas the six-state model explicitly allows each sequence to adopt its own phase and orientation before estimating the shared composition matrix. However, the model assigns each sequence to an inferred offset and orientation state; this state should not be interpreted directly as a biological reading frame. Likewise, the reported synthetic assignment accuracy applies to the specified generating model and should not be interpreted as an accuracy estimate for heterogeneous environmental reads. The single shared composition matrix is a useful description, but heterogeneous samples may contain several distinct triplet-composition patterns.

The practical implications depend on the analytical objective. Bringing sequences into a common inferred phase and randomly redistributing retained offsets answer different questions: the former can emphasize triplet-specific composition, whereas the latter tests whether a read-coordinate pattern depends on phase bias. In particular, reassignment based on the same nucleotide patterns subsequently analyzed can introduce selection effects. We therefore regard phase-aware analysis as a preprocessing framework for specified tasks, rather than a universal use.

Finally, the cross-library survey documents recurrence across multiple studies, rather than estimating prevalence in all ancient or modern metagenomes. Its selected maximum-*R*^2^ summaries depend on the eligible read-length classes and quantify the regularity of the positional pattern, not its absolute amplitude or the fraction of coding reads. With these distinctions maintained, the read-coordinate signal and the latent-state algorithm provide complementary ways to investigate how underlying sequence composition becomes represented relative to fragment boundaries in complex DNA libraries.

## 4 Conclusions

Three-base nucleotide periodicity can remain coherent in read coordinates across complex DNA libraries. The empirical controls, coding-frame analysis, and simulations support an interpretation in which triplet-specific sequence composition interacts with sequence-dependent fragment formation or recovery. Purine-associated boundary selection is sufficient to generate this behavior in the bacterial reference tested, making it a plausible contributor to the signal in ancient environmental libraries without establishing it as the unique cause.

The alignment-free six-state algorithm further showed that, despite distinct read-coordinate profiles across modulo-3 length classes, the inferred triplet-specific nucleotide-composition matrix remained highly stable across read lengths, consistent with a shared underlying compositional structure whose observed phase depends on read boundaries.

The accompanying software provides a reusable framework for investigating these patterns without reference alignment. Validation against biological reading frames and explicit downstream benchmarks will determine when these inferred states can improve sequence analysis. Identification of likely reading frames could support gene prediction and reduce the computational burden of protein-level annotation [29, 30].

Together, these findings motivate distinguishing intrinsic triplet composition from the boundary-associated sampling processes that make it visible in sequencing reads.

## 5 Methods

### 5.1 Datasets and library selection

A2424 (ENA run accession ERR1883436), used as the primary exemplar library for detailed analyses, is an indexed library prepared from sample SP2921 from Denisova Cave (Russia), East Gallery, Layer 11.4 [9]. The library was classified as showing ancient-DNA preservation based on the mammalian mitochondrial-DNA screening criterion used for the source dataset. Ancient environmental DNA libraries were drawn from AncientMetagenomeDir [25]; selected modern examples included agricultural soil [26], Tara Oceans marine metagenome [27], and a human-associated metagenome from the Kawasaki disease study [28]. Reads were stratified by exact observed sequence length. Read-length classes containing at least 40,000 observed reads were eligible for the cross-library analysis. For paired-end datasets, the first read (R1) was used for the read-coordinate composition analysis; single-end datasets were analyzed directly.

### 5.2 Read orientation, filtering, and read-position nucleotide frequencies

Let *ℓ* be the raw read length and Σ = {A, T, G, C}. The analysis pipeline accepts FASTA, FASTQ, BAM, or CRAM. For BAM/CRAM, secondary, supplementary, and quality-control-failed records are excluded. For empirical libraries, identical nucleotide sequences were collapsed within each exact read-length class before nucleotide-frequency profiles were calculated. Each unique sequence therefore contributed once to the positional composition profile. Reads containing symbols outside Σ are excluded before length selection and their number is recorded.

All positions are defined in the original sequenced orientation. For mapped reverse-strand BAM records, the original sequence is restored before analysis using pysam’s forward-sequence accessor; this is necessary because BAM stores the sequence in alignment orientation for reverse alignments. For a retained exact-length class, let *N*_*ℓ*_ denote the number of retained unique, unambiguous nucleotide sequences and let *n*_*b,ℓ*_(*k*) be the number carrying nucleotide *b* at one-based read position *k*. The read-position fraction is

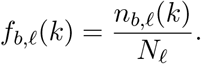

The denominator therefore contains only the same unambiguous reads that contribute to the nucleotide counts. The pipeline exports the complete position-by-base table, plots, per-length counts, excluded-read counts, code hash, Python version, and run parameters.

### 5.3 Period-3 regression, effect size, and permutation testing

The primary regression window is read positions 10–40 inclusive. For the three selected modern metagenomic examples, which contained substantially longer reads and showed stronger terminal composition effects, the common internal window of read positions 30–60 was used. Analysis windows are reported explicitly for each dataset. For each nucleotide and exact read length, we fit

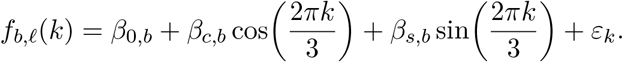

The period-3 coefficient of determination is

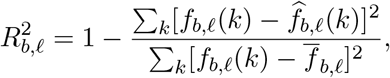

and the fitted oscillation amplitude is

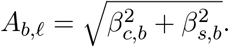

Both quantities are reported because *R*^2^ measures regularity whereas *A*_*b,ℓ*_ measures absolute effect size.

For each nucleotide, a permutation *P* value is obtained by shuffling its observed read-position fractions within the analysis window and refitting the fixed period-3 model. We used 10,000 permutations with deterministic, independent child seeds for A, T, G, and C. The four nucleotide *P* values within each read-length class are adjusted by Holm’s method. These tests do not correct for searching across read lengths; any library-level selection over lengths requires a separate calibrated null procedure.

### 5.4 Random phase redistribution and matched fixed-trim controls

To redistribute read-coordinate phase while preserving sequence composition, A2424 sequences of length 59 were shortened by exactly two nucleotides to a final length of 57 bp. The two removed nucleotides were distributed between the sequenced 5^*′*^ and 3^*′*^ ends using one of three 5^*′*^/3^*′*^ trimming configurations: 0/2, 1/1, or 2/0. These configurations shift the retained 5^*′*^ boundary by zero, one, or two nucleotides and therefore represent the three possible phase offsets modulo three.

For each randomization replicate, a trimming configuration was assigned at the level of the unique original nucleotide sequence, such that all occurrences of an identical sequence received the same assignment. The assignment was deterministic for a given randomization seed and sequence, using a BLAKE2b hash of the seed and original sequence modulo three. Four independent randomization replicates were generated.

As phase-preserving controls, the 0/2, 1/1, and 2/0 configurations were also applied separately and uniformly to all sequences from the same filtered source population. Thus, randomized and fixed-trim datasets were all 57 bp long and differed only in whether multiple retained phase offsets were mixed within the analyzed population. After trimming, identical 57-bp sequences were collapsed, so distinct 59-bp source sequences could occasionally merge and produce small differences in the final unique-sequence counts.

### 5.5 Discrete Fourier representation

The DFT uses read positions 10–39 inclusive, a 30-position window chosen so that frequency 1*/*3 is represented exactly. For nucleotide *b*, let *x*_*b*_(*j*) = *f*_*b,ℓ*_(*k*_*j*_) for *j* = 0, …, *m −* 1 and *m* = 30. We calculate

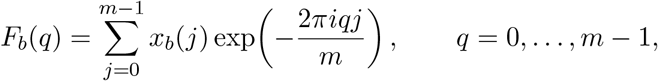

and report the normalized magnitude

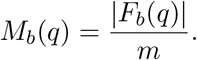

The zero-frequency term is excluded from the displayed spectrum. Period three lies at *q* = 10, *q/m* = 1*/*3. For this non-Nyquist bin, the fraction of the read-position variance carried by the conjugate Fourier pair is

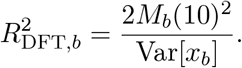

This quantity equals the variance fraction of the corresponding sine–cosine projection up to numerical precision. The implementation uses the explicit base order A–T–G–C and exports the full position-by-base table, spectrum CSV, metadata JSON, PNG, and PDF.

### 5.6 Library-level aggregation

For each library, period-3 regression was performed separately for every eligible read-length class. For each length, a summary score was calculated as the mean of the nucleotide-specific *R*^2^ values for A, T, G, and C. To represent each library in the cross-library analysis, we selected the eligible read length with the largest mean *R*^2^. Each library therefore contributed a single value to the cross-library comparison.

Libraries were grouped according to study/project metadata, and individual library values were displayed as jittered points. The median selected value within each project was used as a descriptive project-level summary. The cross-library analysis was used to assess the breadth and distribution of period-3 structure across independent libraries and projects rather than to perform hypothesis testing on the selected maximum *R*^2^ values.

### 5.7 Six-state hard-EM model

To estimate triplet-specific nucleotide composition while allowing individual sequences to differ in phase and orientation, we implemented a six-state hard expectation–maximization (EM) model [31, 32]. The six latent states correspond to the three possible cyclic offsets in either the forward or reverse-complement orientation.

For each sequence *n* retained at an exact raw length, we construct a 3 *×* 4 nucleotide-count matrix 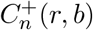 from the sequence and a corresponding matrix 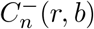 from its reverse complement.

Rows *r ∈ {*0, 1, 2} index sequence positions modulo three and columns index the four nucleotides. The model estimates a row-normalized 3 *×* 4 probability matrix *P* = (*p*_*q,b*_), representing nucleotide probabilities in the three latent position classes.

For orientation *s ∈ {*+, *−}* and cyclic offset *o ∈ {*0, 1, 2}, sequence *n* is assigned the score

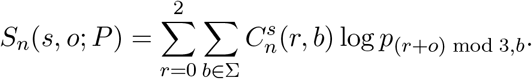

In the hard E-step, each sequence is assigned to the orientation and offset with the largest score. In the M-step, the selected count matrices are realigned according to their assigned offsets, pooled across sequences, and updated with pseudocount *α* before row normalization.

For the empirical A2424 analyses, identical nucleotide sequences were collapsed by exact sequence identity before fitting, so that each distinct sequence contributed once. Each read length was fitted from 20 independently initialized parameter matrices. Each restart iterated the E- and M-steps until the hard assignments were unchanged following parameter re-estimation, with a maximum of 100 iterations. Restarts that did not reach assignment stability within this limit were recorded as nonconvergent and excluded from model selection. Among converged restarts, the solution with the largest complete-data hard-assignment log score was retained. The analysis was considered unsuccessful only if no restart converged.

Because cyclic relabeling of the three position classes and strand-complement transformations generate equivalent representations of the same fitted model, fitted solutions were canonicalized over these label symmetries before comparison across read lengths or datasets.

Model recovery was evaluated using synthetic data. Ten independent positive-control datasets and ten composition-matched IID-null datasets were generated, each containing 60,000 sequences of length 60. Positive-control sequences were generated from a prespecified 3 *×* 4 position-class nucleotide-probability matrix and equally distributed among the six latent phase/orientation states. IID-null sequences were generated using the same overall nucleotide composition but without position-class dependence. Each synthetic dataset was fitted using 20 restarts and a maximum of 500 iterations. Recovery of the latent states was evaluated after resolving the cyclic and strand-complement label symmetries of the model. Because the six states were equally represented in the synthetic positive control, random assignment has an expected accuracy of 1*/*6 (16.7%).

### 5.8 Coding-frame normalization

To test whether the observed three-base periodicity was related to codon-position-dependent sequence composition, we analyzed A2424 reads mapping to annotated coding sequences (CDSs). Read alignments and genomic feature annotations were based on a concatenated reference comprising the Unified Human Gastrointestinal Genome (UHGG) catalog (version 2.0) [33] and the human reference genome GRCh38 (Ensembl release 110) [34]. UHGG reference assemblies and annotations were obtained from MGnify, and the human reference assembly and corresponding GFF3 annotation were obtained from Ensembl. Reads were aligned to the concatenated reference using BWA-MEM v0.7.17-r1188 [35]; alignments were converted to BAM format and coordinate-sorted using SAMtools v1.10 [36].

BAM records were processed using pysam v0.23.3. Primary alignments with mapping quality (MAPQ) *≥* 25 were retained. Reads were required to map unambiguously and in the sense orientation within a single annotated CDS and to be fully contained within that CDS. Alignments with complex CIGAR operations, reads overlapping multiple CDS features, reads containing ambiguous nucleotide symbols, and reads failing the mapping or CDS containment criteria were excluded.

For each accepted read, the annotated CDS strand and GFF3 phase were used to determine its position relative to the coding frame. Zero, one, or two nucleotides were removed from the sequenced 5^*′*^ end as required to place the first retained nucleotide in a common coding-frame class. This produced 69,892 frame-normalized sequences.

For the matched comparison, we selected the 11,094 accepted source reads that produced frame-normalized sequences of length 75. Two FASTA datasets were constructed from exactly the same source reads: a frame-normalized dataset and a matched unnormalized dataset retaining the corresponding sequences without phase-dependent 5^*′*^ normalization. Duplicate collapsing was not applied to this paired comparison so that every accepted source read contributed exactly once to each dataset and the two populations remained matched after transformation. Both datasets were then processed using the same nucleotide-composition and period-3 pipeline, with period-3 regression performed over read positions 10–40.

### 5.9 Simulation of purine-associated fragmentation

To test whether nucleotide-dependent fragmentation could generate coherent read-coordinate period-3 structure, we simulated fixed-length fragments from the *Pseudomonas aeruginosa* PAO1 genome (GenBank accession `AE004091.2`) [37] and the human GRCh38 reference genome (RefSeq assembly `GCF_000001405.26)` [34]. The purine-associated boundary model was motivated by the enrichment of purines at ancient-DNA strand breaks [7, 8, 23].

Candidate fragments were sampled globally across all reference contigs in proportion to the number of valid start positions. Each candidate was assigned to the forward or reverse-complement orientation with equal probability. The read-oriented 5^*′*^ boundary was defined as the nucleotide immediately outside the emitted sequence, whereas the 3^*′*^ boundary was defined as the terminal nucleotide within the emitted sequence.

Purine-associated fragmentation was modeled using a boundary-bias parameter *q ∈* [0.5, 1]. Candidates with A or G at the relevant boundary were accepted with probability one, whereas candidates with C or T were accepted with probability

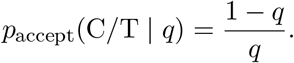

Thus, *q* = 0.5 imposes no nucleotide-dependent boundary selection: valid candidates are accepted irrespective of whether the boundary nucleotide is A, C, G, or T, and the resulting boundary composition reflects the underlying reference sequence. Increasing *q* progressively favors purine-associated boundaries, with *q* = 1 accepting only candidates with A or G at the specified boundary. The bias parameter therefore describes relative boundary acceptance rather than the final purine fraction among accepted fragments.

We evaluated seven 5^*′*^/3^*′*^ conditions: random fragmentation (0.50*/*0.50), 0.55*/*0.50, 0.70*/*0.50, 0.80*/*0.50, 0.90*/*0.50, 1.00*/*0.50, and 1.00*/*1.00. For each reference genome and condition, ten independent replicates of 100,000 reads of length 59 were generated. Candidate sampling and rejection were performed using a seeded deterministic global sampler, and ambiguous candidates were excluded.

Each simulated dataset was analyzed using the same period-3 regression framework. For visualization, the four nucleotide-specific *R*^2^ values were averaged within each replicate. Points in the simulation summary represent the mean of these replicate-level values across the ten independent simulations, and error bars represent 95% *t* confidence intervals across replicate means.

### 5.10 Software and reproducibility

All custom analyses were implemented in Python. Source code is maintained in the project GitHub repository [38]. The exact software release associated with the analyses reported here (version 1.0.0) is permanently archived at Zenodo [39]. The repository includes installation instructions, environment specifications, analysis scripts, validation workflows, and commands required to reproduce the reported analyses.

#### Use of generative artificial intelligence

During manuscript revision, the authors used OpenAI ChatGPT to assist with language editing, organization, and code editing. The authors inspected and revised the output, independently verified the scientific claims and software, and take full responsibility for the manuscript.

## Abbreviations

CDS: coding sequence
DFT: discrete Fourier transform
EM: expectation-maximization
sedaDNA: sedimentary ancient DNA

## Declarations

### Ethics approval and consent to participate

This study performed secondary computational analyses of publicly available sequencing datasets and involved no new recruitment or collection of human samples. The human-associated metagenomic dataset analyzed here was originally collected under a study protocol approved by the institutional medical ethics committees of the University of Tokyo, Tokyo Women’s Medical University, and the National Institute of Infectious Diseases in Japan (Approval No. 295); the original study reports written informed consent from the parents of all participating children [28]. In the present study, no additional identifiable participant information was accessed.

### Consent for publication

Not applicable.

### Availability of data and materials

The primary sequencing datasets analyzed in this study are publicly available from their original repositories. Accession-level provenance for the ancient DNA cross-library analysis, together with the exact AncientMetagenomeDir environmental-library metadata table used for this analysis, is provided with the accompanying software and data release.

Source code is maintained in the project GitHub repository [38] and is released under the MIT License. The exact version of the software, accession-level provenance, metadata, and numerical source data associated with the analyses reported in this study (version 1.0.0) is permanently archived at Zenodo [39].

### Authors’ contributions

S.R., V.S., and M.S. led and supervised the research. All authors contributed to the conceptualization and methodology of the study. N.H. implemented the computational analyses with the support of N.D. and Y.D. All authors contributed to the interpretation of the results and to the review and editing of the manuscript. All authors read and approved the final manuscript.

## Acknowledgements

The authors thank Roded Sharan, David L. Tennenhouse, and Yahli Hecht for taking the time to read the manuscript and for their thoughtful suggestions.

